# Analytical models for *β*-diversity and the power-law scaling of *β*-deviation

**DOI:** 10.1101/2020.04.19.049163

**Authors:** Dingliang Xing, Fangliang He

## Abstract

1. *β*-diversity is a primary biodiversity pattern for inferring community assembly. A randomized null model that generates a standardized *β*-deviation has been widely used for this purpose. However, the null model has been much debated and its application is limited to abundance data.
2. Here we derive analytical models for *β*-diversity to address the debate, clarify the interpretation, and extend the application to occurrence data.
3. The analytical analyses show unambiguously that the standardized *β*-deviation is a quantification of the effect size of non-random spatial distribution of species on *β*-diversity for a given species abundance distribution. It robustly scales with sampling effort following a power law with exponent of 0.5. This scaling relationship offers a simple method for comparing *β*-diversity of communities of different sizes.
4. Assuming logseries distribution for the metacommunity species abundance distribution, our model allows for calculation of the standardized *β*-deviation using occurrence data plus a datum on the total abundance.
5. Our theoretical model justifies and generalizes the use of the *β* null model for inferring community assembly rules.

## INTRODUCTION

Comparing *β*-diversity, the variation in species composition, among different regions is an important yet controversial topic in ecology (Kraft et al., 2011; Qian, Chen, Mao, & Ouyang, 2013; Bennett & Gilbert, 2016; Ulrich et al., 2017). A randomization-based null model that was said to correct for the dependence of raw *β*-diversity on species pool (Kraft et al., 2011) has been widely adopted for this purpose (e.g., De Cáceres et al., 2012; Myers et al., 2013; Vannette & Fukami, 2017; Xing & He, 2019; Zhang, He, Zhang, Zhao, & von Gadow, 2020). However, this randomization approach has been criticized on several aspects, including incorrect interpretations of the *β*-deviation metric and the dependence of the metric on sampling effort and species pool (Qian et al., 2013; Bennett & Gilbert, 2016; Ulrich et al., 2017, 2018). Another limitation of the null model is that it requires individual-level data and thus limits its application to situations when only abundances of individual species in each local community are available. Therefore, there is a strong need for a theoretical scrutiny of the null model to justify its use and also a need to extend its application to presence/absence data.

It has been established that the abundance of species and their spatial aggregation in a metacommunity are the two basic quantities that construct macroecological patterns of diversity, including *β*-diversity (He & Legendre, 2002; Plotkin & Muller-Landau, 2002; Morlon et al., 2008). All ecological processes, biotic and abiotic, are acting through these two quantities to affect the observed macroecological patterns (He & Legendre, 2002). For instance, both dispersal limitation and environmental filtering can result in spatially aggregated distribution of species, which in turn contributes to high *β*-diversity. However, the reverse is not necessarily true: a high *β*-diversity does not have to result from aggregated distribution. This is because differences in species pool could also alter values of observed *β*-diversity. Therefore, it is necessary to disentangle the effect of spatial aggregation from that of species pool for inferring contributions from different ecological processes on *β*-diversity.

Different from the original *β*-diversity, Kraft et al.’s null model (and the resultant standardized *β*-deviation, or *β*-deviation for short) was designed to preserve species abundance distribution (SAD) of metacommunity while randomizing the species-level spatial distribution. It is thus clear that the *β*-deviation is a measure of the effect of non-random spatial distribution of species on *β*-diversity for a given SAD. Although this link between *β*-deviation and intraspecific aggregation of species was acknowledged in previous studies (e.g., Crist, Veech, Gering, & Summerville, 2003; Myers et al., 2013), including Kraft et al. (2011), this interpretation of *β*-deviation has not been widely heeded in the literature (e.g., Qian et al. 2013). The original interpretation of *β*-deviation was “a standard effect size of *β*-diversity deviations from a null model that corrects for γ dependence” (Kraft et al., 2011) but this misinterpretation attracted much criticism on the null model (Qian et al., 2013; Bennett & Gilbert, 2016; Ulrich et al., 2017). To reiterate, *β*-deviation is neither a measure that corrects for the effect of γ diversity nor a measure to correct for the effect of SAD (Xu, Chen, Liu, & Ma, 2015). While these debates stimulated searches for alternative null models aiming to correct for the γ-dependence (Ulrich et al., 2018), the problem of the misinterpretation has not been solved but undermines the use of otherwise a promising and important approach in community ecology (Myers et al., 2013; Vannette & Fukami, 2017; Xing & He, 2019). Here by developing an analytical null model, we reinforce that *β*-deviation measures the effect of species spatial distribution on *β*-diversity.

A second problem of the *β*-deviation is that the metric is subject to the effect of sampling effort (Bennett & Gilbert, 2016). In a recent study, we demonstrated by simulation that *β*-deviation can effectively identify and correctly compare non-random *β* patterns across assemblages under the condition of constant sampling effort (see Appendix S2 in Xing & He 2019). This suggests that there might be some scaling relationship between *β*-deviation and the sampling effort. In this study, we show that there is actually a surprisingly simple and robust scaling relationship between *β*-deviation and sampling effort. The discovery of this scaling relationship is remarkable. It analytically addresses the criticisms raised by Bennett & Gilbert (2016) on the sample-effort dependence of *β*-deviation.

The last shortcoming of the currently used null model, inherited from the randomization procedure, is that it only deals with abundance data. Yet, in many studies, particularly at regional scales, presence/absence of species are the only data available (e.g., Gaston et al., 2007; McKnight et al., 2007; Buckley & Jetz, 2008). Therefore, there is a need to extend its application to presence/absence data, so that it could be used in broader contexts such as delineating biogeographical regions (Kreft & Jetz, 2010).

Here, based on theories about species abundance distribution and spatial distribution of species, we develop an analytical null model to replace the randomization procedure and to address all the above problems. The analytical results reveal that Kraft et al.’s *β*-deviation (1) is a measure of the effect size of non-random spatial species distribution on *β*-diversity, (2) scales with the sampling effort following a power law with a constant exponent of 0.5, and (3) can be reasonably estimated based on presence/absence data, provided that a datum on the total abundance of the metacommunity is known (which is usually more readily available than abundance for each species). We test the performance and utility of our analytically derived models using several well-studied empirical data sets.

### DERIVATION OF ANALYTICAL MODELS FOR *β*-DIVERSITY

Consider a metacommunity consisting of *M* equal-sized (equal-area) local communities and there are in total *N* individuals belonging to *S* species. The widely used proportional species turnover (denoted *β*_P_ following Tuomisto (2010)) is defined as 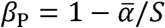, where 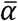 is the average number of species over the *M* local communities. Two other major multi-community measures of *β*-diversity, namely the *M*-community Sorensen-(*β*_S_) and Jaccard-differentiation (*β*_J_) (Chao & Chiu, 2016), are monotonically related with 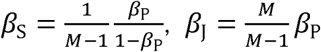. Hence, although we will only focus on *β*_P_ in this paper, our analytical models can be easily transformed to these other indices.

From the definition and some simple rearrangements (see Appendix S1), it is easy to show that *β*_P_ is the expectation of the probability that a species from the metacommunity is absent from a local community and thus can be expressed as:

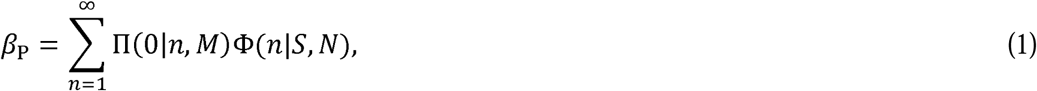

where Π(0|*n, M*) is the probability that a species with *n* individuals in the metacommunity is absent from a randomly chosen local community, and Φ(*n*|*S, N*) is the metacommunity species abundance distribution (SAD), i.e., the probability of a species having *n* individuals. This model provides a general framework to develop analytical solutions for Kraft et al.’s null model and the resultant *β*-deviation.

### Null model: *β*-diversity under random spatial distribution of species

Kraft et al.’s null model randomly shuffles species identities across the *N* individuals that comprise the *M* local communities while keeping the metacommunity SAD and each of the local community sizes unchanged. By repeating the randomization procedure a large number of times (e.g., 999) one can get the expectation (*β*_null_) and corresponding variance (Var_Π_(*β*_null_)) of *β*-diversity under the null model. Here the subscript Π is used to denote that the variation is due to species spatial distribution, in contrast to that due to SAD, which will be made clear below. This randomization is equivalent to assuming random spatial patterns of species. *β*-deviation is then defined as 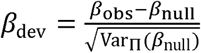 (Kraft et al. 2011). This standardized metric describes the degree of departure from random distribution in spatial distribution of empirical species, while maintaining the SAD Φ(*n*|*S, N*).

We now use the general formulation of *β*_P_ as defined by equation (1) to derive the null *β* for random distribution of species in the metacommunity consisting of *M* local communities. The probability for a randomly distributed species with abundance *n* being absent from a local community is: 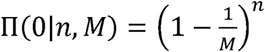. What we need to determine is which SAD should be used in equation (1). The empirical SAD was used in Kraft et al. (2011). It is well established in theory that SAD of metacommunity follows logseries distribution. This is a principal result of the neutral theory of ecology (Hubbell, 2001; Volkov, Banavar, Hubbell, & Maritan, 2003) and is also predicted by the maximum entropy theory of ecology (Pueyo, He, & Zillio, 2007; Harte, 2011). In practice, logseries together with lognormal distribution have been repeatedly shown to be the two models fitting empirical data best (McGill et al., 2007) and recent meta-analyses even show logseries outperforms other models (White, Thibault, & Xiao, 2012; Baldridge, Harris, Xiao, & White, 2016). In this study, we adopt the theoretical logseries SAD for metacommunity to derive *β*_null_ and its variance. We also tested the use of lognormal distribution by simulation but that does not change our main results about the scaling of *β*-deviation (see Appendix S2).

For a logseries SAD, 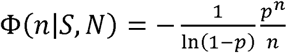, the only parameter *p* = *e* ^− *λ*^ is fully determined by the state variables *N* and 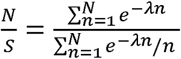 (Harte, 2011). Based on this SAD model and the binomial model for random species spatial distribution (Barton & David, 1959; He & Reed, 2006), we derive the following analytical version of the null model (see Appendix S1 for mathematical details):

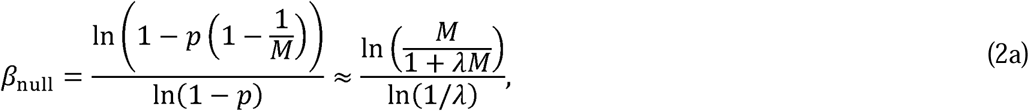

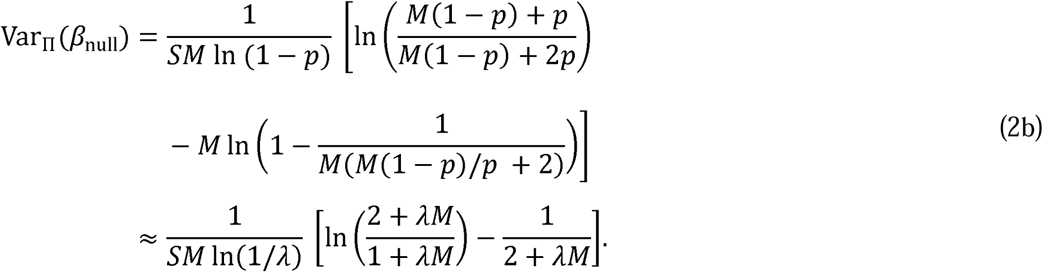

We note that the approximations are valid under conditions M ≫ 1 and *λ* ≪ 1 which are expected for real ecosystems (Harte 2011, p150). This is because in real applications the size of a local community is usually much smaller than the area of the metacommunity (*M* = metacommunity area / local-community size). However, we would recommend using the exact formulas in real applications, particularly when the sample size is small, since they impose no computing challenge. However, the approximations offer analytical simplicity and we will use them to derive the scaling relationship between *β*-deviation and the sampling effort (see below).

From equation (2) it is clear that the expectation and variance of *β*_null_ under the spatial random distribution are fully determined by the parameter of the logseries SAD (*p*), the number of local communities (*M*), and the total number of species (*S*). The property that the parameter *p* is fully determined by *S* and *N* allows *β*_null_ to be parameterized using only the three state variables *N, S*, and *M*. This means we can compare *β*-diversity calculated from occurrence data with the null model as long as the metacommunity size *N* is known (or can be estimated). This offers a method of testing *β*-diversity in cases where abundances of individual species are not available. The resultant analytical *β*-deviation is obtained by substituting equation (2) into 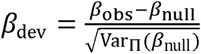.

The variance in equation (2b) is the expected conditional variance due to species spatial distribution, i.e., Var_Π_ (*β*_null_) = E_Φ_ (Var_Π_ (*β*| Φ)). No variation due to SAD arises because in Kraft et al.’s randomization approach the empirical SAD is used. However, in scientific inference, the empirical SAD ought to be considered as a sample of the underlying theoretical logseries SAD. It then becomes clear that the sampling procedure introduces additional variance in the expected *β*_null_. According to the law of total variance, this second variance can be written as (Appendix S1): 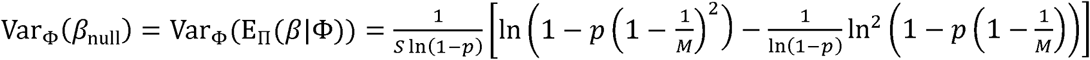, where the subscript Φ indicates that the variation is due to SAD. This leads naturally to a quantification of the variance of *β*-deviation due to the metacommunity SAD:

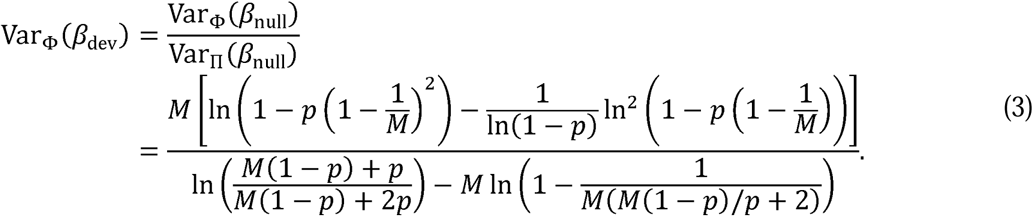

### *β*-diversity under aggregated spatial distribution of species

In reality, species almost always show aggregated instead of random spatial distribution due to a variety of ecological processes such as dispersal limitation, habitat filtering, and local competition. It has long been recognized that spatial aggregation of empirical species can be well modeled by the negative binomial distribution (NBD, Boswell & Patil 1970; Pielou 1977). Using the same logseries SAD model as before and the NBD for aggregated species spatial distribution, we can derive a prediction of *β*_P_ (Barton & David 1959; He & Reed 2006; see Appendix S1 for the derivation):

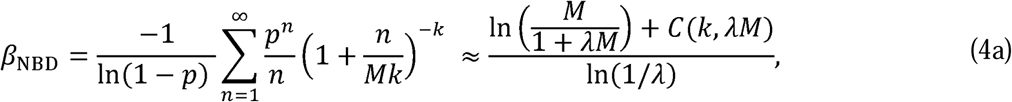

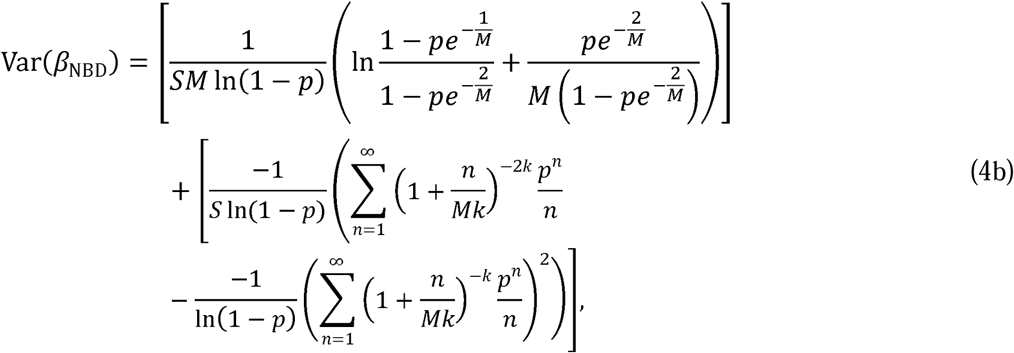

where k (> 0) is an aggregation parameter with smaller values indicating more aggregated distribution, *C*(*k, λM*) is a function fully determined by *k* and *λM* (i.e., a constant for given values of *k* and *λM*; see Appendix S1). Like equation (2), the approximation here is valid under conditions *M* ≫ 1 and *λ* ≪ 1. Note that the variance in equation (4b) includes both the variation due to species spatial distribution (the first square bracket) and the variation due to SAD (the second square bracket). This is different from the null model (2), where only the variance due to random species spatial distribution occurs.

Model (4) provides a framework to predict *β*-diversity from the three state variables *N, S*, and *M* and the information about species spatial aggregation (*k*). Different from the null *β* of model (2), model (4) incorporates realistic spatial distribution of species and is expected to describe *β*-diversity of empirical communities (see EMPIRICAL TEST below for confirmation). The realism of model (4) allows us to explore various properties of *β*-diversity in relation to spatial distribution of species. As an example, below we derive a scaling relationship between *β*-deviation and sampling effort, to address a major criticism of the using of *β*-deviation.

### Scaling of *β*-deviation with sampling effort

Substitute the approximate versions of equations (2) and (4a) into Kraft et al.’s 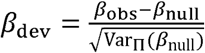 (replace *β*_obs_ with *β*_NED_), we have:

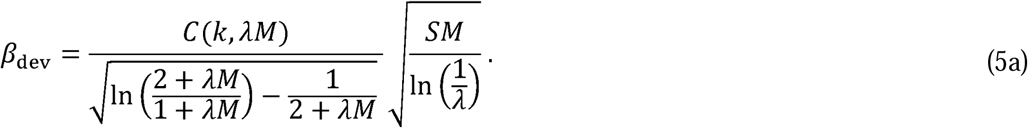

Until now all the derivations in the above assume that the metacommunity is fully sampled. That is, all the *M* local communities (that comprise the metacommunity) are sampled. In real applications we can only sample a fraction of a metacommunity and sampling intensity also changes from study to study. This sampling incompleteness can have consequences. As shown in Bennet & Gilbert (2016), *β*-deviation is subject to the effect of sampling effort. Considering equation (5a), when only *m* (out of *M*) local communities are sampled, the *k* will not change but both *S*(*m*) and *λ*(*m*) will change with *m*. Nevertheless, numerical analyses suggest that *λ*(*m*)*m* and *S*(*m*) (1 / *λ*(*m*)) are nearly constant if parameter *M* ≫ 1 (see also Harte 2011, p243). Hence, for a sample of *m* local communities, the complicated term 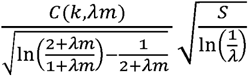 in equation (5a) becomes a constant independent of *m*. The relationship between *β*-deviation and the sampling effort (*m*) turns out to be a simple power law with scaling exponent of 0.5:

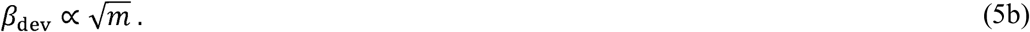

## EMPIRICAL TEST OF THE MODELS

### Data

We used several well-studied data sets from the literature to evaluate the analytical results derived above. Alwyn Gentry’s forest transect data (Phillips & Miller, 2002) were used to confirm the performance of our analytical models (2) and (4). This data set has been previously analyzed using the *β*-deviation (Kraft et al., 2011; Qian et al., 2013). The data consist of 226 plots distributed worldwide, of which 198 are ‘standard’ plots. Each plot is considered as a metacommunity (Kraft et al., 2011; Qian et al., 2013) that contains ten 2×50 m transects (each transect is considered as a local community) where all plants with a stem diameter ≥ 2.5 cm were measured and identified to species or morphospecies. Following the previous studies, we used the 198 standard plots in this study. For each metacommunity, we calculated the observed *β*-diversity (*β*_obs_) and the expected *β*-diversity (*β*_null_), variance Var_Π_ (*β*_null_), and *β*-deviation (*β*_dev_) for the spatial random hypothesis (equation 2). We also computed *β*_dev_ following the randomization approach of Kraft et al. (2011). We then plotted these two *β*_dev_ to evaluate consistency between our analytical null model and the original randomization approach. These two *β*_dev_ were also plotted versus absolute latitude for comparison. Our analytical models assumed logseries SAD for the metacommunities. To test this assumption and to evaluate potential consequences due to violations of this assumption, a Kolmogorov-Smirnov goodness-of-fit test was performed for each of the metacommunities.

With respect to the NBD model (equation 4), we tested its performance by plotting the predicted versus observed *β*-diversities. Beyond the three state variables (i.e., *N, S*, and *M*) that are known for each metacommunity, model (4) requires an additional aggregation parameter *k*. We estimated it for each metacommunity by maximizing the log-likelihood function (He, Gaston, & Wu, 2002): 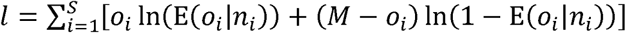 for the occupancy model 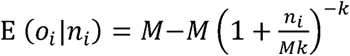, where *o*_*i*_ is the number of local communities where a species with *n*_*i*_ individuals is present (i.e., the occupancy). The other notations are the same as in previous sections.

To test the prediction about the power law scaling of *β*-deviation with sampling effort, we extracted data on *β*-deviation and sample size from the four different datasets reported in Bennet & Gilbert (2016). The four datasets are: (1) plants in 605 1-m^2^ plots collected from meadow patches in the Garry Oak Ecosystem of southern British Columbia and northern Washington State; (2) plants in 110 1-m^2^ plots collected in an abandoned field at the Koffler Scientific Reserve in southern Ontario, Canada; (3) understory plants in 85 50-m^2^ forest plots collected from Mount St. Hilaire, near Montreal, Canada; and (4) diatoms in surficial sediments sampled from 492 North American lakes. More details about these datasets can be found in Bennet & Gilbert (2016) and the references therein. The extracted *β*-deviation and sample size were plotted on a log-log graph and were compared against the theoretical power law of equation (5b).

### Test results

The empirical tests shown in Fig. 1 confirm that our theoretical null model (equation 2) for the random species spatial distribution accurately implements the randomization procedure of Kraft et al. (2011). The analytical null model *β*-deviation has an appreciable but weaker correlation with the *β*-deviation computed from Kraft et al.’s randomization process (*R*^2^ > 0.49, Fig. 2a). Only nine out of Gentry’s 198 metacommunities show significant difference between the two versions of *β*-deviation. Both methods are consistent in showing the decreasing latitudinal gradients of *β*-deviation, but the gradient is appreciably somewhat stronger using the analytical *β*-deviation (Fig. 2b, c). The Kolmogorov-Smirnov test shows that the logseries describes the metacommunity SAD very well (*P* > 0.05) for 180 out of the 198 plots (Fig. 3). All the nine “outlier” plots identified in Fig. 2 are among the few that have a poor fit of logseries (*P* < 0.05; Fig. 3b).

**Figure 1.**
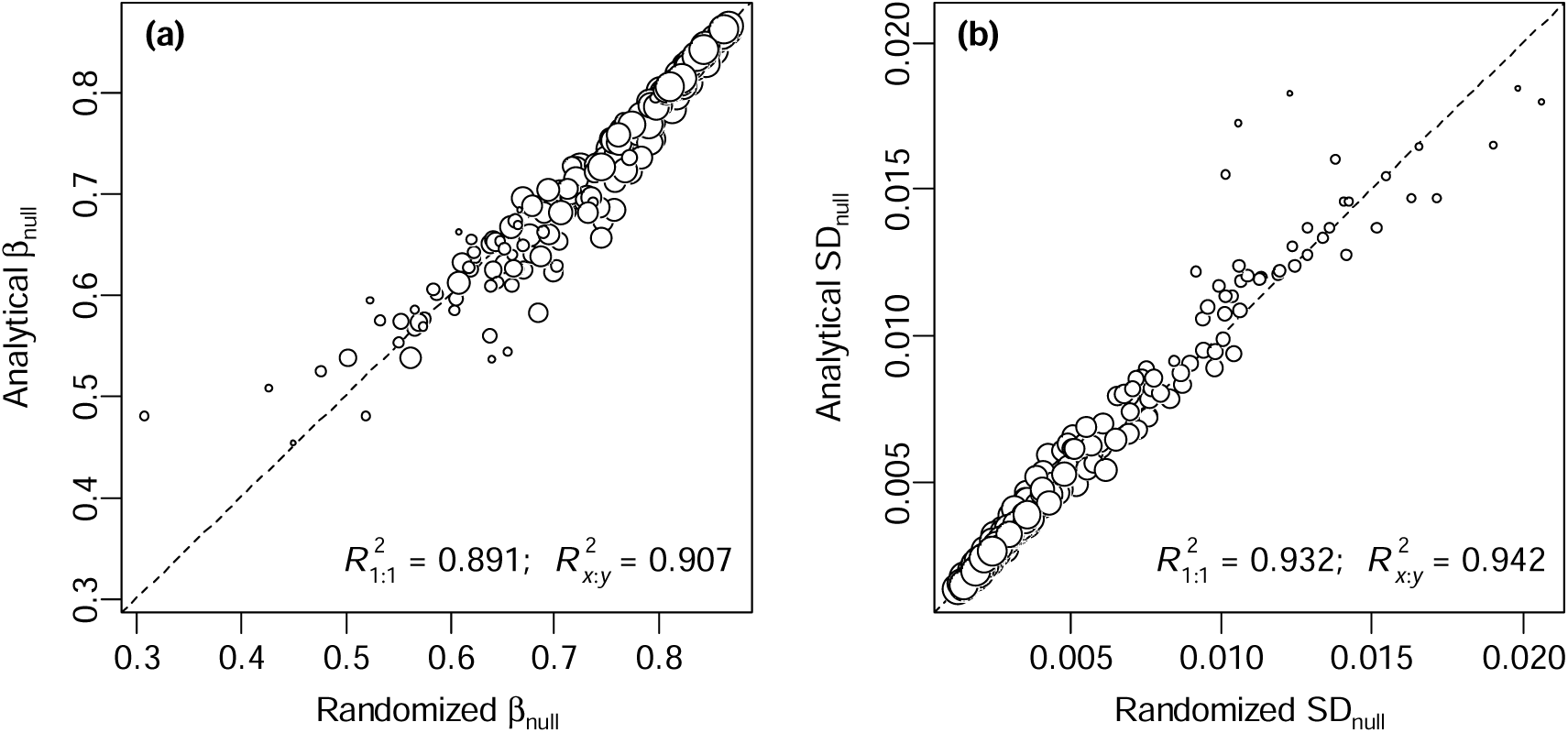
*β*-diversity of Gentry’s 198 forest plots under the null model of random spatial distribution of species. (a) Relationship between the expected *β*-diversity calculated using equation (2a) and the randomization approach of Kraft et al. (2011). (b) Relationship between standard deviation of *β*_null_ calculated using equation (2b) and the randomization approach. Sizes of the points are proportional to the values of logarithm of the number of species. The dashed lines are 1:1 line 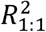 describes the goodness-of-fit of the 1:1 line to the data, while 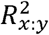 is the *R*^2^ between *x* and *y*.

**Figure 2.**
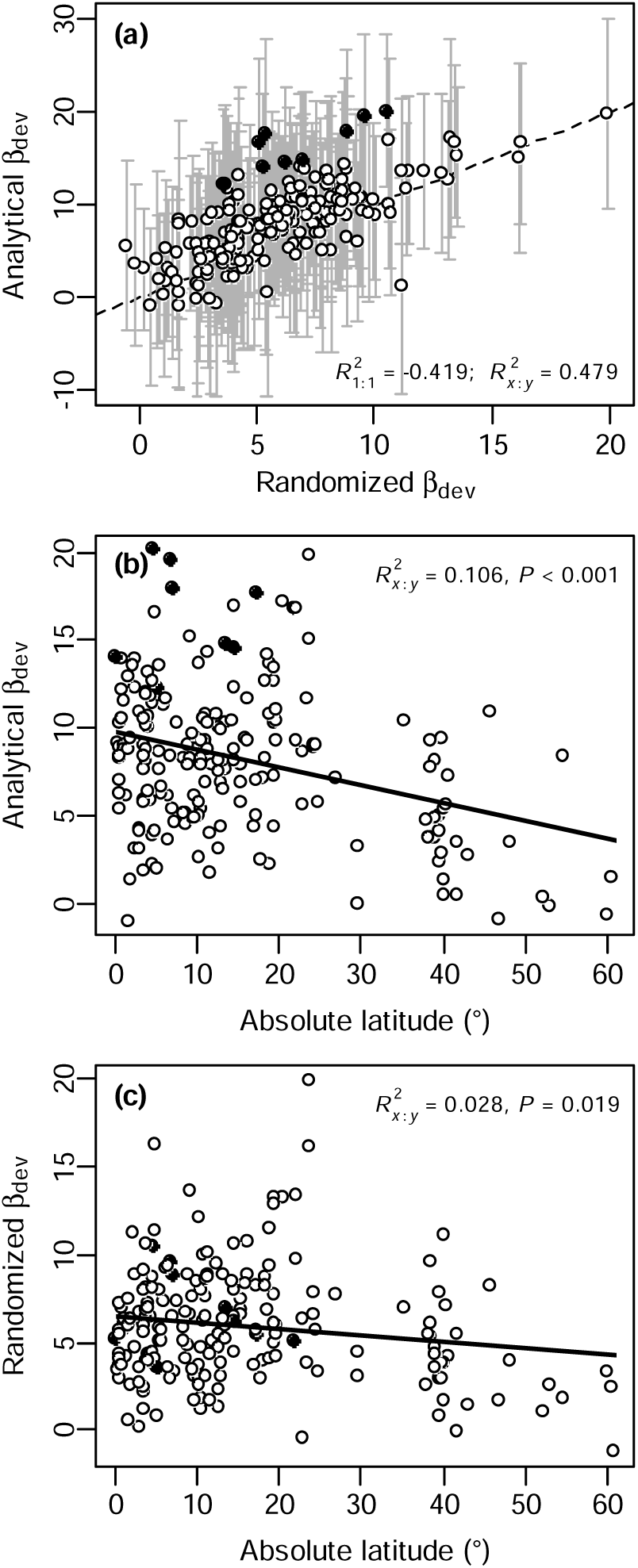
*β*-deviation for Gentry’s 198 forest plots. (a) Relationship between *β*-deviation calculated using the analytical equation (2) and the randomization approach. Error bars represent 1.96 SD of *β*-deviation due to species abundance distribution (equation 3). The dashed line is 1:1 line. 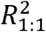 describes how well the 1:1 line fits the data. (b) Relationship of *β*-deviation with latitude computed using the analytical solution, and (c) the randomization approach. The solid lines in (b, c) show ordinary least squares regression. 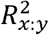 is the *R*^2^ between *x* and *y*. Filled points highlight the nine plots where the two *β*-deviations are significantly different.

**Figure 3.**
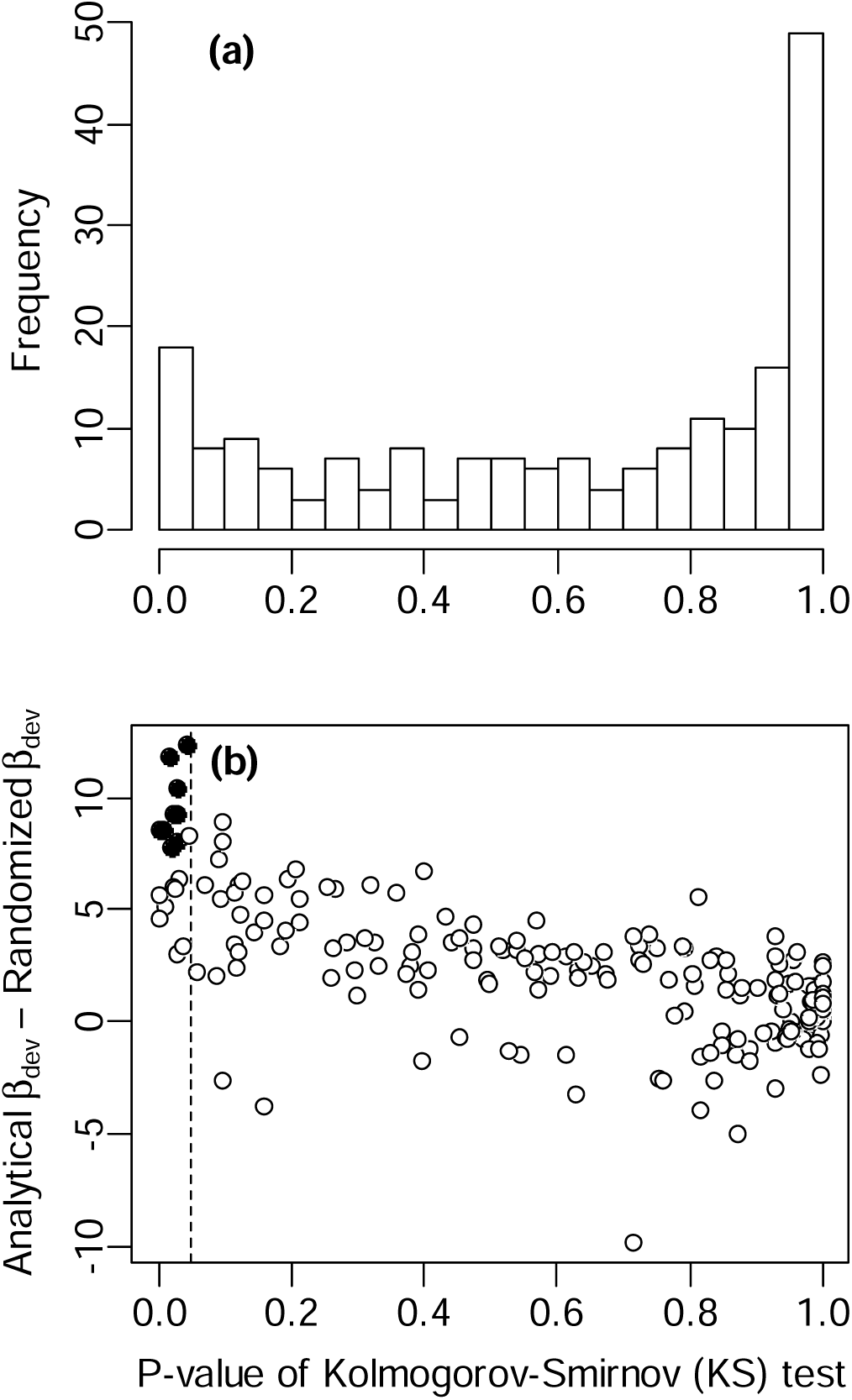
Goodness-of-fits of the logseries SAD to Gentry’s data. (a) Histogram of *P*-value of the Kolmogorov-Smirnov (KS) test for the logseries SAD model. (b) Relationship of the difference between the analytical and randomization *β*-deviation with *P*-value of the KS test. Filled points in (b) are plots where the two *β*-deviations are significantly different (same as in Figure 2). The dashed vertical line indicates *P* = 0.05.

The NBD *β*-diversity (equation 4) performs very well in predicting empirical *β*-diversity (*R*^2^ = 0.88, Fig. 4). Again, nine out of Gentry’s 198 metacommunities show significant departure from the prediction (note seven of those nine are the same “outlier” plots as in Fig. 2). The empirical relationships between *β*-deviation and sampling effort employed by Bennett & Gilbert (2014) to argue against the use of *β*-deviation follow our theoretical power-law scaling equation (5b) very well, as long as the sample size is not too small (i.e., *m* > 30, Fig. 5).

**Figure 4.**
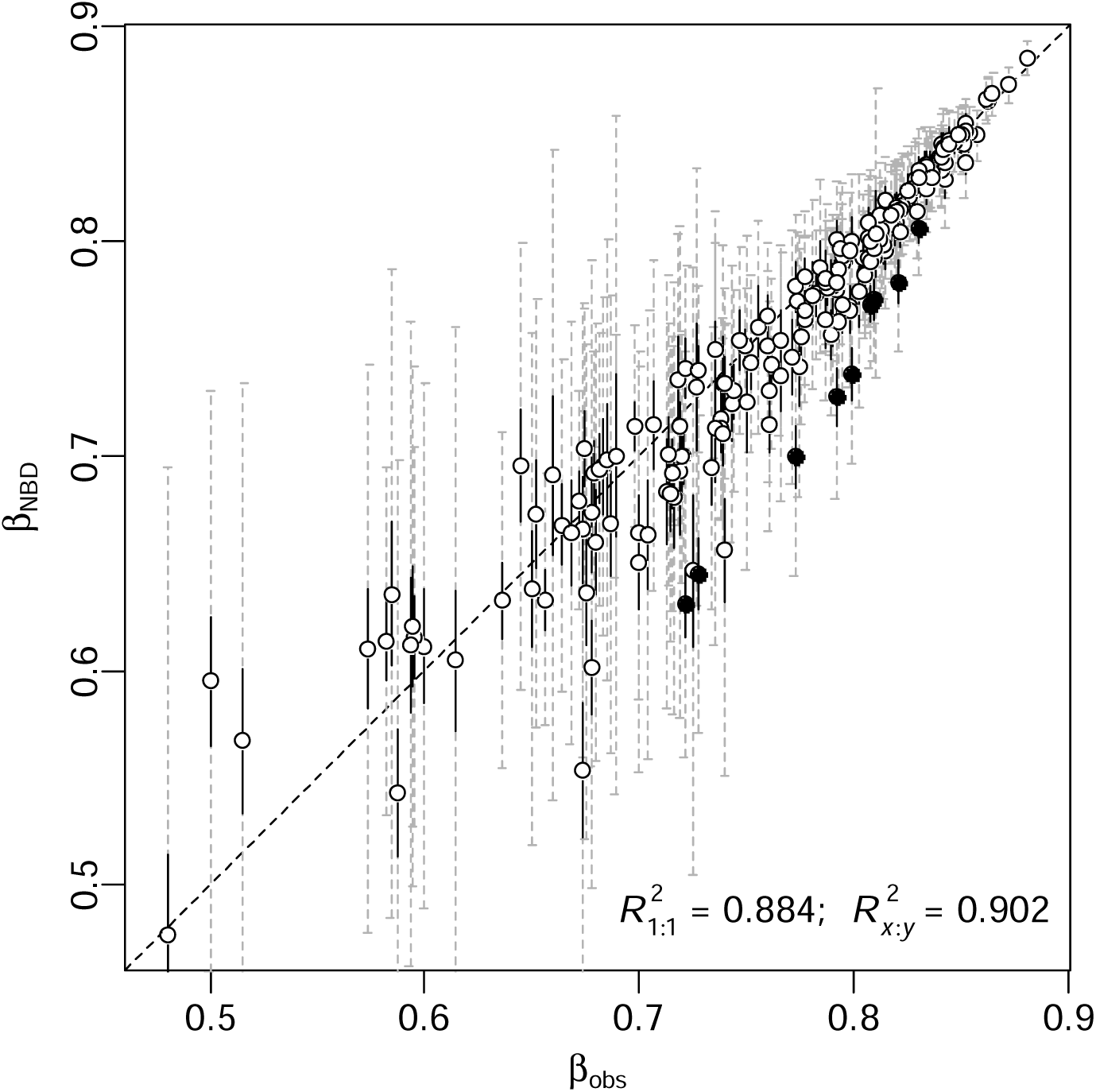
Relationship between predicted *β*-diversity using model (4) and observed *β*-diversity for Gentry’s 198 forest plots. The grey dashed error bars represent 1.96 SD due to both species abundance distribution and species-level spatial distribution, with the black solid portions due to species-level spatial distribution only. The dashed line is the 1:1 line. Filled points represent the nine plots where the observed *β*-diversity differ significantly from the prediction. 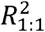 describes the goodness-of-fit of the 1:1 line to the data, while 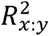 is the *R*^2^ between the two *β*-diversity.

**Figure 5.**
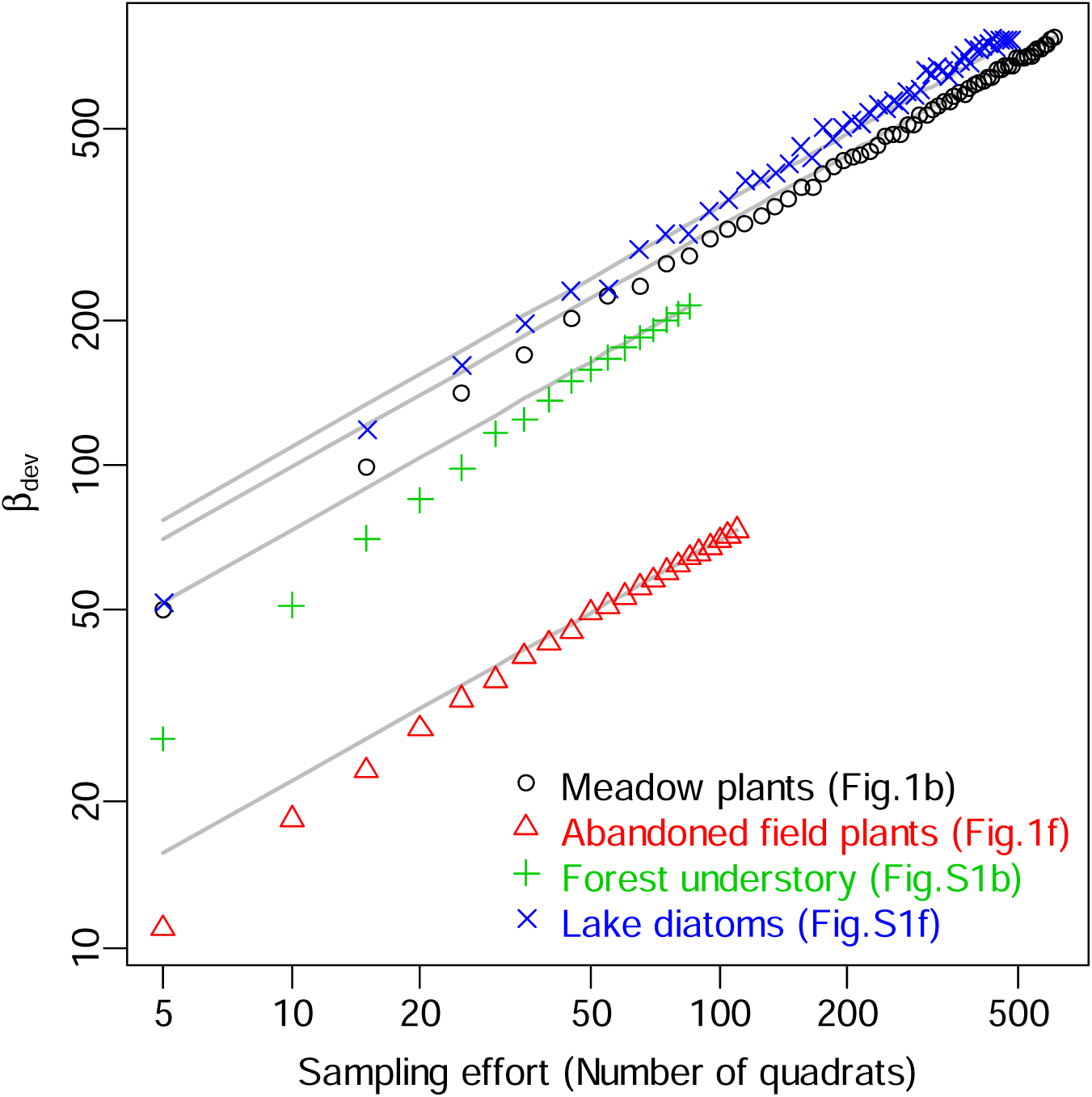
Relationships of *β*-deviation with sampling effort for the four different datasets from Bennett & Gilbert (2016). The grey lines are the predictions of the derived power-law with scaling exponent of 0.5 (equation 5b). The figure numbers in the legend refer to figures in Bennett & Gilbert (2016) where the data were extracted.

## DISCUSSION

In this paper we have developed analytical *β*-diversity for random and aggregated spatial distribution. The random *β*-diversity provides a null model for standardizing empirical *β*-diversity which would otherwise has to be implemented by randomization (Kraft et al., 2011). The simulation and empirical tests show that the analytical *β*-diversity models perform very well in comparison to the original randomization process of Kraft et al. (2011) (Figs. 1-2). The practical difference between the theoretical null model and the original randomization approach can be simply explained by the use of different SADs, which is one of two differences between our null model and the original randomization approach. The second difference is that we do not preserve the abundance in each local community. Previous study has demonstrated that this second difference is neglectable (see Fig. 3a in Xu et al., 2015). Our null model uses a theoretical distribution (could be logseries, or lognormal; see Appendix S2), while the randomization approach uses the empirical SAD that could be subject to sample size and sampling variation. The variation arising from sampling SAD is quantified by equation (3). Recognizing the existence of this variation is important because in real applications almost all empirical SADs are based on sampling data and themselves are not true metacommunities. This sampling problem is one of the major criticisms of Kraft et al.’s (2011) original analyses of Gentry’s data that the empirical SAD does not represent true metacommunity (Tuomisto & Ruokolainen, 2012). From this point of view, our analytical *β*-diversity based on the theoretically justified logseries SAD of metacommunity provides a desirable solution.

Unlike the randomization process, our *β*-diversity models do not require abundances of individual species but only the data on metacommunity size (*N*) and total richness (*S*) (for null *β*-diversity equation 2) or *N* and *S* plus spatial distribution parameter *k* (for the NBD *β*-diversity equation 4). More significantly, these analytical results reveal the dependence of *β*-diversity on *N, S*, species spatial pattern and sampling effort and thus offer unambiguous interpretation on *β*-deviation which is otherwise not obvious and has been a subject of debate (Kraft et al., 2011; Qian et al., 2013; Bennett & Gilbert, 2016; Ulrich et al., 2017). In the following discussion, we will address and clarify the controversies surrounding the use of the randomized *β*-deviation. Our discussion will make it clear that *β*-deviation is an important and solid diversity measure for testing community assembly mechanisms and the much-criticized sampling effect on *β*-deviation (Bennett & Gilbert, 2016) can be easily dealt with using the simple scaling law of equation (5).

The fact that the null *β*-diversity of equation (2) is derived from the assumption of random spatial distribution of species clearly indicates that the randomization procedure of Kraft et al. (2011) is for randomizing spatial distribution of species, not for correcting for the effect of regional (γ) diversity on *β*-diversity as originally interpreted. Hence, the *β*-deviation is a metric measuring the effect size of non-random species spatial distribution on *β*-diversity for a given SAD.

The misinterpretation of *β*-deviation as “a standard effect size of *β*-diversity deviations from a null model that corrects for γ dependence” (Kraft et al., 2011) has raised much concern about the null model and led to attempts to develop alternative null models or metrics to truly correct for “γ dependence” (Qian et al., 2013; Ulrich et al., 2017, 2018). It is clear from equation (5a) that *β*-deviation for a metacommunity with a fixed number of local communities (*M*; can also be thought as a fixed sampling design) is fully determined by a non-linear, interactive function of the two parameters *k* and *λ* for the metacommunity (Fig. S1 in Appendix S1). It is also known that the parameter *λ* is a function of the ratio between total abundance *N* and the γ-diversity *S* (Harte, 2011). Given the well-known non-linear relationship between *N* and *S*, it is inevitable for *β*-deviation to be dependent on *S* (imagine two systems having the same *M* and *k*: to maintain a constant *β*-deviation they must have the same *λ*, which requires the same *N*/*S* ratio, which can only be true for the same *S* value). This further clarifies that *β*-deviation is not correcting for the “γ dependence”.

There are a number of alternative approaches used in the literature that are closely related with the null model investigated here, but they all suffer from similar problems. Qian et al. (2013) and Xu et al. (2015) analyzed the raw (i.e., *β*_obs_ – *β*_null_), instead of the standardized, *β*-deviation. From our models (2) and (4), it is clear that the raw *β*-deviation is also dependent on γ, thus not serving as a correction for the γ-dependence. A more fundamental problem with the raw *β*-deviation is that it is not a measure of effect size and thus is of little use for comparing the effect of species aggregation on *β*-diversity. Ulrich et al. (2018) proposed a “fixed-fixed” null model that preserves both row and column sums of the community matrix to correct for the γ-dependence. However, the “fixed-fixed” null model does not have an analytical solution and its ecological interpretation is not clear.

Another major criticism on *β*-deviation is that it is sampling-effort dependent, meaning that different sample sizes from the same system can lead to drastically different *β*-deviation (Bennett & Gilbert 2016). This problem makes the method not useful for comparing different communities unless sample size is standardized (Xing & He, 2019; Zhang et al., 2020). The scaling relationship between *β*-deviation and sampling effort (equation 5b) derived in this study thaws this criticism and offers a simple solution. To compare *β*-deviation with different sample size (*m*), one only needs to standardize the effect of sample size by dividing the *β*-deviation by 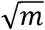. This analytical result is a generalization of the simulation study of Xing & He (2019) (see Appendix S2 there). It is remarkable because the simulation study only justifies comparisons of *β*-deviation under constant sampling effort. It does not provide the standardization method revealed by the analytical study presented here. The standardization method revealed here is critical for future comparison study or meta-analyses of published results from different studies. However, it is important to note that in applications the scaling relationship does not behave as well for small sample sizes (e.g., *m* < 30).

The null model of *β*-diversity and the resultant *β*-deviation provide a promising approach for inferring community assembly rules. The derivation of the analytical version of the null model in this study clarifies the interpretation of the model and addresses the problem that the *β*-deviation is subject to the effect of sampling effort. The requirement for species abundance data by the original randomization null model can seriously limit its application in systems of large spatial extent. Data on abundance of individual species are no longer needed for the theoretical model derived here as long as the size of the metacommunity is known. This affords the use of the theoretical model to systems of large extent and can take advantage of the occurrence data available for many taxa (e.g., birds, amphibians, mammals, and plants; Gaston et al., 2007; McKnight et al., 2007; Buckley & Jetz, 2008; Maitner et al., 2018). Coupling with these data, our analytical model offers a possibility to explore the global pattern and underlying drivers of *β* diversity.

One potential limitation of this study is the assumption of logseries for metacommunity SAD. In situations where the true SAD does not follow logseries, our model could elevate type I error (i.e., the *β*-deviation tends to reject a true hypothesis that communities are randomly assembled in space). That is, deviation of the true SAD from logseries will weaken the accuracy of predictions from our models (Fig. 3). As for the power-law scaling of *β*-deviation with sampling effort, both our simulation (Appendix S2) and the empirical data from various sources (Fig. 5) show that it is robust to variations in SAD. The ecological interpretation of *β*-deviation revealed by our analytical analyses does not suffer from the SAD assumption either. Hence, among the three major implications of our analytical results, only the application in estimating *β*-deviation from occurrence data could potentially be affected by the assumption. Because logseries for metacommunity SAD is theoretically justified (Hubbell, 2001; Pueyo et al., 2007; Harte, 2011) and well supported in numerous empirical tests (White et al., 2012; Baldridge et al., 2016), our theoretical models are of strong practical significance, as exemplified by the majority of Gentry’s forest plots (Figs. 1-4). However, erring on the side of caution, we suggest avoiding using our model if the SAD of the study system is far from logseries.

## ACKNOWLEDGEMENTS

We thank late Dr. Alwyn H. Gentry and his co-workers for collecting the invaluable forest transect data set and thank the Missouri Botanical Garden for making the data publicly available. DX thanks Prof. John Harte for generously sending a copy of his book (Harte, 2011). This work was supported by the Natural Science and Engineering Research Council of Canada.

## Notes

https://github.com/DLXING/beta_analytical

### Competing Interest Statement

The authors have declared no competing interest.

