## Appendix S1 for "Analytical models for *β*-diversity and the power-law scaling of *β*-deviation"

**Appendix S1. Derivation of the analytical models for beta diversity**

**Derivation of equation (1)**

For an $S\times M$ metacommunity presence/absence matrix, denote $\alpha_{i} \{i=1,\cdots,M\}$ as the number of species in the *i*-th local community (i.e., the sum of the *i*-th column of the matrix), $o_{j} \{j=1,\cdots,S\}$ as the number of occurrences of the *j*-th species (i.e., the sum of the *j*-th row of the matrix). Then $\Pi\left( 0 | n_{j},M \right)=1-\frac{o_{j}}{M}$ is the probability that the *j*-th species (with *n_j_* individuals in the metacommunity) is absent from a randomly chosen local community. The following derivation is straightforward:

$$\beta_{P}=1-\frac{\bar{\alpha}}{S}$$

$$=1-\frac{\sum_{i=1}^{M} \frac{\alpha_{i}}{M}}{S}$$

$$=1-\frac{\sum_{j=1}^{S} \frac{o_{j}}{M}}{S}$$

$$=\frac{\sum_{j=1}^{S} \left( 1-\frac{o_{j}}{M} \right)}{S}$$

$$=\text{E}_{\Phi}\left( \Pi\left( 0 | n,M \right) \right)$$

$$=\sum_{n=1}^{\infty} \Pi\left( 0 | n,M \right)\Phi(n|S,N).$$

**Derivation of equation (2)**

Under the maximum entropy theory of ecology (METE), it has been shown that species-abundance distribution (SAD) of metacommunity follows logseries distribution (Pueyo *et al*. 2007; Harte 2011):

$\Phi\left( n | S,N \right)=-\frac{1}{\ln\left( 1-p \right)}\frac{p^{n}}{n}$.

It is worth to note that under neutral theory SAD of metacommunity also follows the logseries distribution (Hubbell 2001; Volkov *et al*. 2003). Let’s now randomly sample an individual from the logseries SAD and place that individual to one of the *M* local communities. The probability that a species with abundance *n* is absent from a local community is: $\Pi\left( 0 | n,M \right)=\left( 1-\frac{1}{M} \right)^{n}$. (This is equivalent to assuming that species in the metacommunity are randomly distributed among the *M* local communities.) Substituting this absence probability and the logseries SAD to equation (1), we have equation (2a):

$$\beta_{\text{null}}=\sum_{n=1}^{\infty} \left( 1-\frac{1}{M} \right)^{n}\frac{-1}{\ln\left( 1-p \right)}\frac{p^{n}}{n}$$

$$=\frac{\ln\left( 1-p\left( 1-\frac{1}{M} \right) \right)}{\ln(1-p)}.$$

To derive the expected conditional variance of *β*_null_ due to species spatial distribution, we use $\beta=\frac{\sum_{j=1}^{S} \left( 1-\frac{o_{j}}{M} \right)}{S}$ from the above derivation and the variance of $o_{j}$ under random spatial distribution (Barton & David 1959; He & Reed 2006): $\mathrm{Var}\left( o_{j} | n_{j} \right)=M\left( M-1 \right)\left( 1-\frac{2}{M} \right)^{n_{j}}+M\left( 1-\frac{1}{M} \right)^{n_{j}}-M^{2}\left( 1-\frac{1}{M} \right)^{2n_{j}}$. Based on this information, we have equation (2b):

$$\text{Var}_{\Pi}\left( \beta_{\text{null}} \right)=E_{\Phi} \left( \text{Var}_{\Pi}\left( \beta| \Phi\right) \right)$$

$$=E_{\Phi}\left( \left( \frac{1}{SM} \right)^{2}\sum_{j=1}^{S} \text{Var}_{\Pi}\left( o_{j} | n_{j} \right) \right)$$

$$=E_{\Phi}\left( \left( \frac{1}{SM} \right)^{2}\sum_{j=1}^{S} \left( M\left( M-1 \right)\left( 1-\frac{2}{M} \right)^{n_{j}}+M\left( 1-\frac{1}{M} \right)^{n_{j}}-M^{2}\left( 1-\frac{1}{M} \right)^{2n_{j}} \right) \right)$$

$$=\frac{1}{SM}E_{\Phi}\left( \frac{1}{S}\sum_{j=1}^{S} \left( \left( M-1 \right)\left( 1-\frac{2}{M} \right)^{n_{j}}+\left( 1-\frac{1}{M} \right)^{n_{j}}-M\left( 1-\frac{1}{M} \right)^{2n_{j}} \right) \right)$$

$$=\frac{1}{SM}\sum_{n=1}^{\infty} \left( \left( M-1 \right)\left( 1-\frac{2}{M} \right)^{n}+\left( 1-\frac{1}{M} \right)^{n}-M\left( 1-\frac{1}{M} \right)^{2n} \right)\frac{-1}{\ln\left( 1-p \right)}\frac{p^{n}}{n}$$

$$=\frac{1}{SM\ln\left( 1-p \right)} \left[ \left( M-1 \right)\ln\left( 1-p\left( 1-\frac{2}{M} \right) \right)+\ln\left( 1-p\left( 1-\frac{1}{M} \right) \right)-M\ln\left( 1-p\left( 1-\frac{1}{M} \right)^{2} \right) \right]$$

$$=\frac{1}{SM ln(1-p)} \left[ \ln\left( \frac{M\left( 1-p \right)+p}{M\left( 1-p \right)+2p} \right)-M\ln\left( 1-\frac{1}{M\left( M\left( 1-p \right)/p +2 \right)} \right) \right].$$

**Derivation of equation (3)**

Based on the definition of *β*-deviation: $\beta_{\text{dev}}=\frac{\beta_{\mathrm{obs}}-\beta_{\mathrm{null}}}{\sqrt{\text{Var}_{\Pi}(\beta_{\text{null}})}}$, its variance due to SAD is simply $\mathrm{Var}_{\Phi}\left( \beta_{\text{d}\text{ev}} \right)=\frac{\mathrm{Var}_{\Phi}\left( \beta_{\text{null}} \right)}{\mathrm{Var}_{\Pi}\left( \beta_{\text{null}} \right)}$, where $\mathrm{Var}_{\Phi}\left( \beta_{\text{null}} \right)$ is the variation of *β*_null_ due to SAD. So equation (3) follows from the derivation of $\mathrm{Var}_{\Phi}\left( \beta_{\text{null}} \right)$ below:

$$\text{Var}_{\Phi}\left( \beta_{\text{null}} \right)=\mathrm{Var}_{\Phi}\left( E_{\Pi}\left( \beta| \Phi\right) \right)$$

$$=\mathrm{Va}r_{\Phi}\left( \frac{\sum_{j=1}^{S} \left( 1-\frac{1}{M} \right)^{n_{j}}}{S} \right)$$

$$=\frac{1}{S}\mathrm{Va}r_{\Phi}\left( \left( 1-\frac{1}{M} \right)^{n} \right)$$

$$=\frac{-1}{S\ln\left( 1-p \right)}\left( \sum_{n=1}^{\infty} \left( 1-\frac{1}{M} \right)^{2n}\frac{p^{n}}{n}-\frac{-1}{\ln\left( 1-p \right)}\left( \sum_{n=1}^{\infty} \left( 1-\frac{1}{M} \right)^{n}\frac{p^{n}}{n} \right)^{2} \right)$$

$$=\frac{1}{S\ln\left( 1-p \right)}\left[ \ln\left( 1-p\left( 1-\frac{1}{M} \right)^{2} \right)-\frac{1}{\ln\left( 1-p \right)}\ln^{2} \left( 1-p\left( 1-\frac{1}{M} \right) \right) \right].$$

**Derivation of equation (4)**

We use negative binomial (NBD) to model aggregated spatial distribution. Under NBD, we have (Barton & David 1959; He & Reed 2006): $\Pi\left( 0 | n,M \right)=\left( 1+\frac{n}{Mk} \right)^{-k}$and $\text{Var}\left( o_{j} | n_{j} \right)=Me^{-\frac{2n_{j}}{M}}\left( e^{\frac{n_{j}}{M}}-1-\frac{n_{j}}{M} \right)$. So like the above derivations, we can easily derive equation (4a) for expected *β*-diversity:

$$\beta_{\mathrm{NBD}}=\frac{-1}{\ln\left( 1-p \right)}\sum_{n=1}^{\infty} \frac{p^{n}}{n}\left( 1+\frac{n}{Mk} \right)^{-k}$$

$$\approx\frac{\ln\left( \frac{M}{1+\lambda M} \right)+C\left( k, \lambda M \right)}{\ln\left( \frac{1}{\lambda} \right)},$$

and equation (4b) for the total variance of the expectation:

$$\mathrm{Var} \left( \beta_{\mathrm{NBD}} \right)=E_{\Phi} \left( \text{Var}_{\Pi}\left( \beta| \Phi\right) \right)+\text{Var}_{\Phi}\left( \text{E}_{\Pi}\left( \beta| \Phi\right) \right)$$

$$=\left[ \frac{1}{SM\ln\left( 1-p \right)}\left( \ln\frac{1-pe^{-\frac{1}{M}}}{1-pe^{-\frac{2}{M}}}+\frac{pe^{-\frac{2}{M}}}{M\left( 1-pe^{-\frac{2}{M}} \right)} \right) \right]+\left[ \frac{-1}{S\ln(1-p)}\left( \sum_{n=1}^{\infty} \left( 1+\frac{n}{Mk} \right)^{-2k}\frac{p^{n}}{n}-\frac{-1}{\ln\left( 1-p \right)}\left( \sum_{n=1}^{\infty} \left( 1+\frac{n}{Mk} \right)^{-k}\frac{p^{n}}{n} \right)^{2} \right) \right].$$

The validity of the approximate version of equation (4a) is demonstrated in Fig. S1 below.

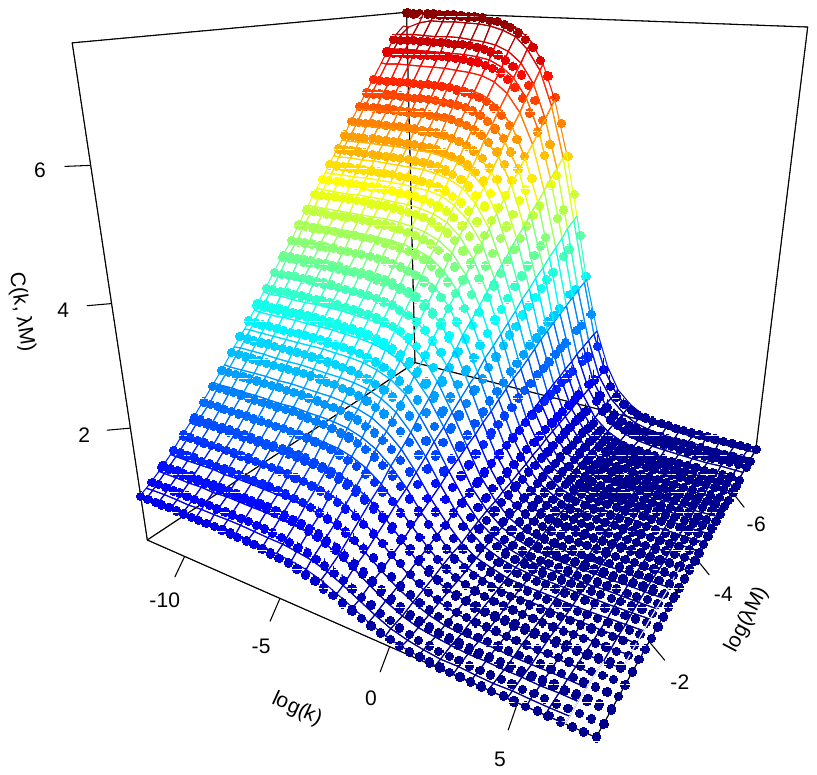

**Figure S1** A numerical test illustrating that $\sum_{n=1}^{\infty} \left[ \left( 1+\frac{n}{Mk} \right)^{-k}\frac{e^{-\lambda n}}{n} \right]-\ln\left( \frac{M}{1+\lambda M} \right)$ is fully determined by *k* and *λM*. The dots represent values calculated using the exact expression. The surface is the relationship fitted from the numerical analysis and the fitted relationship is $C(k,\lambda M)=\frac{\ln\left( \frac{1}{\lambda M}+1 \right)}{1+\left( 5.07-0.44\ln\left( \lambda M \right) \right)k}$.
