## Appendix S2 for "Analytical models for *β*-diversity and the power-law scaling of *β*-deviation"

**Appendix S2. Simulation results justifying the validity of the analytical models for communities having lognormal species abundance distributions**

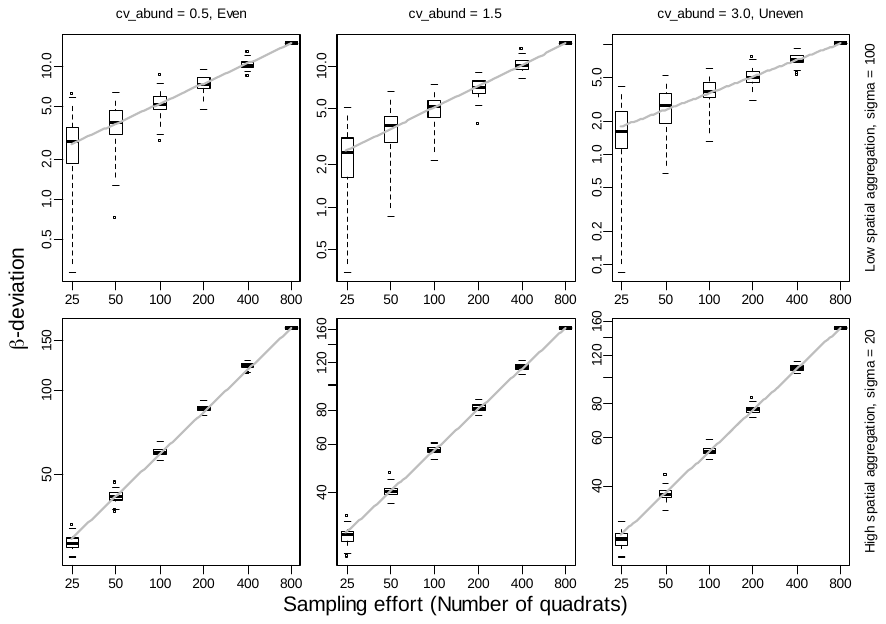

**Figure S2** Relations of *β*-deviation with sampling effort for six simulated communities. The communities were simulated using the sim_thomas_community() function of the mobsim R package (version 0.1.0). Species abundance distribution (SAD) was set to follow lognormal distribution by controlling the sad_type argument. All communities had the same spatial extent (500×1000), number of species (100), and total abundance (20,000), but they had different SAD evenness (controlled by the cv_abund argument; indicated above the 1^st^ row of the panels), and different degree of intraspecific aggregation (controlled by the sigma argument; indicated on the right side of the 2^nd^ column of the panels). Each community was divided into 800 quadrats of size 25×25 and *β*-deviation was calculated for samples of varying sampling effort (the *x*-axis) using the randomization procedure. Each boxplot represents distribution of results from 100 replications of the sampling. The grey lines represent the predicted power-law scaling with an exponent of 0.5.
